# Altered neocortical gene expression, brain overgrowth and functional over-connectivity in *Chd8* haploinsufficient mice

**DOI:** 10.1101/143552

**Authors:** Philipp Suetterlin, Shaun Hurley, Conor Mohan, Kimberley L. H. Riegman, Marco Pagani, Angela Caruso, Jacob Ellegood, Alberto Galbusera, Ivan Crespo-Enriquez, Caterina Michetti, Yohan Yee, Robert Ellingford, Olivier Brock, Alessio Delogu, Philippa Francis-West, Jason P. Lerch, Maria Luisa Scattoni, Alessandro Gozzi, Cathy Fernandes, M. Albert Basson

## Abstract

Truncating *CHD8* mutations are amongst the highest confidence risk factors for autism spectrum disorders (ASD) identified to date. Here, we report that *Chd8* heterozygous mice display increased brain size, motor delay, hypertelorism, pronounced hypoactivity and anomalous responses to social stimuli. Whereas gene expression in the neocortex is only mildly affected at mid-gestation, over 600 genes are differentially expressed in the early postnatal neocortex. Genes involved in cell adhesion and axon guidance are particularly prominent amongst the down-regulated transcripts. Resting-state functional MRI identified increased synchronised activity in cortico-hippocampal and auditory-parietal networks in *Chd8* heterozygous mutant mice, implicating altered connectivity as a potential mechanism underlying the behavioural phenotypes. Together, these data suggest that altered brain growth and diminished expression of important neurodevelopmental genes that regulate long-range brain wiring are followed by distinctive anomalies in functional brain connectivity in *Chd8^+/-^* mice. Human imaging studies have reported altered functional connectivity in ASD patients, with long-range under-connectivity seemingly more frequent. Our data suggest that *CHD8* haploinsufficiency represents a specific subtype of ASD where neuropsychiatric symptoms are underpinned by long-range over-connectivity.

## Introduction

Autism spectrum disorder (ASD) is diagnosed on the basis of socio-communicative deficits and repetitive, perseverative behaviours with restricted interests (APA 2013). ASD is frequently associated with comorbidities like hyper-sensitivity to sensory stimuli, seizures and anxiety (Croen et al. 2015; Jeste and Tuchman 2015; Tavassoli et al. 2014). The phenotypic and genetic heterogeneity of ASD has hampered the elucidation of the molecular mechanisms that may underlie specific aberrant behaviours. However, the recent identification of de novo, likely gene disrupting (LGD) mutations that show highly significant associations with autism (Iossifov et al. 2014; Neale et al. 2012; O’Roak et al. 2014; O’Roak et al. 2012b; Talkowski et al. 2012) provides an opportunity to phenotype and molecularly characterise genetically defined ASD subtypes.

Exome sequencing studies of several thousand simplex families detected de novo, LGD mutations in the *CHD8* (Chromodomain Helicase DNA binding factor 8) gene (Iossifov et al. 2014; Neale et al. 2012; O’Roak et al. 2014; O’Roak et al. 2012b; Talkowski et al. 2012). Patients with *CHD8* mutations are characterised by a high incidence of autism, macrocephaly, facial dysmorphisms, motor delay and hypotonia, intellectual disability and gastro-intestinal problems (Bernier et al. 2014; Merner et al. 2016; Stessman et al. 2017; Stolerman et al. 2016). *CHD8* encodes an ATP-dependent chromatin remodelling protein of the chromodomain helicase DNA binding family (Thompson et al. 2008; Yuan et al. 2007). The recruitment of CHD8 to gene promoters in mouse and human neural progenitors is strongly associated with transcriptional activition, while *CHD8* knock-down in these cells results in the reduced expression of many ASD-associated genes (Cotney et al. 2015; Sugathan et al. 2014).

Three groups recently described *Chd8^+/-^* mouse models (Gompers et al. 2017; Katayama et al. 2016; Platt et al. 2017). Megalencephaly, subtle but wide-spread transcriptional changes and behavioural anomalies were found in all these *Chd8^+/-^* mouse lines. Individual studies reported attenuated expression of neural genes and de-repression of REST (Katayama et al. 2016), alterations in striatal neurotransmission (Platt et al. 2017) and a developmental RNA splicing phenotype (Gompers et al. 2017). Understanding the contribution of each of these mechanisms to the ASD phenotype remains a major challenge.

Altered brain connectivity, characterised by local over-connectivity and long-range underconnectivity, has been hypothesised to underpin some of the neuropsychiatric phenotypes observed in ASD (Belmonte et al. 2004; Just et al. 2004). Resting-state functional MRI (rsfMRI) studies in ASD patients have provided evidence for reduced long-range synchronisation in spontaneous brain activity (reviewed in Picci et al. 2016). Increased long-range connectivity has also been reported in a subset of cases (Di Martino et al. 2014), consistent with the phenotypic heterogeneity of ASD. Thus, the exact nature of aberrant functional connectivity in ASD may depend on the specific underlying aetiology.

Similar rsfMRI studies in ASD mouse models may help bridge the gap between ASD models and the human condition (Liska and Gozzi 2016). As one example, homozygous *Cntnap2* mouse mutants exhibit hypo-connectivity of the default mode network (Liska et al. 2017), a phenotype often observed in idiopathic ASD patients (Cherkassky et al. 2006) and recapitulating analogous clinical observations in humans with *CNTNAP2* mutations (Scott-Van Zeeland et al. 2010).

In the present study we generated a novel *Chd8^+/-^* mouse model. We report behavioural anomalies, macrocephaly and functional over-connectivity in cortico-hippocampal networks in these mice that are prefigured by dysregulation of the cortical transcriptome in the early postnatal period.

## METHODS

### *Chd8* gene targeting

A 14.84kb genomic DNA fragment was subcloned from C57BL/6 BAC clone (RP23: 318M20) into pSP72 (Promega). This fragment encompassed a 9.45kb 5’ long homology arm (LA) and a 4.4kb 3’ short homology arm (SA). The targeting construct was generated by inserting a loxP/FRT-PGK-gb2-Neo cassette 214bp 3’ of exon 3 (inGenious Targeting Laboratory (iTL), Ronkonkoma, NY, USA). An additional single loxP site containing a BclI restriction site for Southern blot screening was inserted 5’ of exon 3. The final targeting construct of 18.8 kb was linearised by NotI digestion and electroporated into C57BL/6J ES cells. G418-resistent clones were selected, screened by PCR and Southern blot for successful homologous recombination. Five clones with successful recombination were identified (Supplementary Fig. 1) and two clones (124 and 254) were injected into Balb/c blastocysts (iTL). Resulting chimaeras were bred with Flpe deleter mice on a C57BL/6J background to excise the neo cassette and produce *Chd8^flox/+^* mice (Supplementary Fig. 1). *Chd8^flox/+^* mice were then crossed with *β-actinCre* mice (Lewandoski and Martin 1997) to generate a *Chd8* null allele *(Chd8^-^). β-actinCre;Chd8^+/-^* mice were crossed with C57BL/6J mice to remove the Cre transgene and establish a *Chd8^+/-^* line.

### Mice

Experimental mice were produced by *Chd8^+/-^* × C57BL/6J crosses, taking care to equalise paternal or maternal inheritance of the *Chd8* null allele, especially for behavioural experiments. For genotyping, genomic DNA was extracted using Proteinase K digestion or the HotSHOT method (Truett et al. 2000). Genotyping reactions were then performed for the presence of *Chd8* wildtype and null alleles using the following primer pair: <u>FW</u>: CCC ACA TCA AGT GGC TGT AA, <u>Rev</u>: GGT AGG GAA GCA GTG TCC AG.

This yielded a PCR product of 395bp for the null allele and 1.1kb for the wildtype allele.

### Western Blot

Telencephalic vesicles were dissected from E12.5 embryos and total cell protein prepared by lysing in 8M urea, 1% CHAPS, 50mM Tris (pH 7.9) containing protease inhibitors (PMSF, Pepstatin A, Leupeptin, Aprotinin; Roche) and a phosphatase inhibitor cocktail (Sigma). Samples were loaded (10μg total protein per lane) onto a Mini-PROTEAN pre-cast gel (BioRad) and resolved using gel electrophoresis. Protein was transferred to a nitrocellulose membrane (Bio-Rad) which was then blocked in 5% non-fat milk powder (Bio-Rad) and 1% bovine serum albumin (BSA, Sigma) in TBS with 0.1% Tween-20 (TBST), followed by incubation with anti-CHD8 primary antibody (rabbit anti-Chd8 N-terminal, Bethyl Laboratories (cat#: A301-224A), 1:5000) in 3% non-fat milk powder and 1% BSA in TBST overnight at 4°C. After washing, the membrane was incubated with HRP-conjugated secondary antibody (Millipore), HRP detected with Clarity ECL reagent (Bio-Rad) and the membrane imaged using a Bio-Rad ChemiDoc system. The same membrane was subsequently incubated with anti-GAPDH primary antibody (rabbit anti-GAPDH, Abcam (cat#: ab9485), 1:40000) overnight at 4°C and probed with HRP-conjugate and imaged as before. Relative protein quantity was calculated using Bio-Rad ImageLab software.

### X-ray Computed tomography

Fixed heads from adult (26 – 27 days old) *Chd8^+/-^* and *Chd8^+/+^* mice (n=7 of each from two different litters) were scanned using a GE Locus SP microCT scanner. The specimens were immobilised using cotton gauze and scanned to produce 28μm voxel size volumes, using a X-ray tube voltage of 80kVp and a tube current of 80μA. An aluminium filter (0.05mm) was used to adjust the energy distribution of the X-ray source. Reconstructions of computer tomography scans, images and measurements were done in MicroView 2.5.0 software (Parallax Innovations, ON, Canada). Each 3D landmark point was recorded, twice for each sample, using the 3D point recording built-in tool within the same software, with the operator blind to the genotypes. The distances between the landmarks were normalised for each sample to the average of the wild-type littermates. Graphics of the plotted data and statistical analysis were performed using GraphPad Prism version 6.0h for Mac OS X (GraphPad Software, La Jolla California USA, www.graphpad.com). Unpaired student t-tests were applied to analyse the variation between the two groups, for every distance between 2 specific 3D landmark points. Three-dimensional coordinate locations of a total of 22 biological relevant cranial landmarks were chosen based on a landmark list for adult mouse skull (Hill et al. 2009).

### Behavioural assessments

Mice for behavioural testing were housed, marked for identification and behaviours assessed essentially as described in Whittaker et al. (2017). Different batches of mice were used for (i) recording pup USVs and spontaneous motor behaviours, and (ii) adult behaviours (9-12 weeks of age at the start of testing; 19-22 weeks of age at the end of testing). For adult behaviours, tests were carried out in the following order: rotarod, grip strength, open field, self-grooming, marble burying, adult social investigation, 3 chamber social approach, light/dark test, olfactory habituation/dishabituation and Morris water maze.

#### General activity measurements

General activity was measured using a running wheel paradigm. Mice were housed individually under a 12h:12h light-dark cycle (lights on at 8am; lights off at 8pm) in a light-, air-, temperature-controlled ventilated cabinet (Arrowmight, Hereford, UK). Running-wheel cages were equipped with an infrared sensor (Bilaney consultant Ltd, Sevenoaks, UK) connected to a computer. Data were collected in 1-min bins using Clocklab software (Actimetrics, Inc, Wilmette, IL, USA). Mice were continuously monitored undisturbed from the day they were placed in the running wheel cages and their general activity during the light versus dark phase were compared over the first 7 days.

### Structural MRI

After completion of adult behavioural tests, mice were terminally anesthetized and intracardially perfused. Samples were processed, imaged and analysed as previously described (Whittaker et al. 2017).

### Resting-State fMRI

rsfMRI experiments were performed on 15-18 week old mice (n=23 *Chd8^+/+^*; n=19 *Chd8^+/-^).* Animals were prepared for imaging as previously described (Ferrari et al. 2012; Sforazzini et al. 2016). Briefly, mice were anaesthetised using isoflurane (5% induction), intubated and artificially ventilated (2% maintenance). Blood pressure was monitored continuously by cannulating the left femoral artery, also allowing for terminal arterial blood sampling. Administration of isoflurane was ceased after surgery and substituted with halothane (0.75%). 45 mins after isoflurane cessation functional data acquisition commenced. To rule out possible genotype-dependent differences in anesthesia sensitivity we continuously recorded two independent readouts previously shown to be linearly correlated with anesthesia depth: arterial blood pressure and amplitude of cortical BOLD signal fluctuations (Liu et al. 2011; Steffey et al. 2003; Zhan et al. 2014). Arterial blood pressure (p=0.79; Supplementary Fig. 5A) and the amplitude of BOLD signal fluctuations of motor cortex (p=0.56; Supplementary Fig. 5B) did not significantly differ between *Chd8^+/-^* mice and littermate controls, eliminating a confounding contribution of anaesthesia to the observed functional hyper-connectivity. In vivo images were obtained using a 7.0 T MRI scanner (Bruker Biospin, Milan), as previously described (Liska et al. 2016). Signal transmission and reception were achieved using a 72mm birdcage transmit coil and a 4-channel solenoid coil. For each session, high resolution anatomical images were acquired using a fast spin echo sequence based on the following parameters: repetition time (TR)/echo time (TE) 5500/60ms, matrix 192 × 192, field of view 2 × 2cm^3^, 24 coronal slices, and slice thickness 0.5mm. Cocentred BOLD rsfMRI time series were acquired using an echo planar imaging (EPI) sequence with the following parameters: TR/TE 1200/15ms, flip angle 30°, matrix 100 × 100, field of views 2 × 2cm^2^, 24 coronal slices, slice thickness 0.5mm, 500 volumes and 10min total acquisition time. Raw MRI data, templates and code employed to generate functional maps are available by contacting AG.

#### Functional Connectivity Analyses

To allow for T_1_ equilibration effects, the first 20 volumes of rsfMRI data were removed. The time series were then despiked, corrected for motion and spatially normalised to an in-house mouse brain template (Sforazzini et al. 2014). Normalised data had a spatial resolution of 0.1042 × 0.1042 × 0.5mm^3^ (192 × 192 × 24 matrix). Mean ventricular signal (averaged rsfMRI time course within a reference ventricular mask) and head motion traces were regressed out of each time series. No genotype-dependent differences were observed in ventricular volume, as measured by the dimensions of individual ventricular masks. All rsfMRI time series were then spatially smoothed (full width at half maximum of 0.6mm) and band-pass filtered using a frequency window of 0.01-0.1Hz.

To identify brain regions displaying genotype-dependent differences in functional connectivity in an unbiased manner, we calculated global rsfMRI connectivity maps for all subjects, as described previously in detail (Liska et al. 2017; Liska et al. 2015). A previously described seed-based approach was then used to examine between-group differences in the intensity and scope of long-range rsfMRI correlation networks (Sforazzini et al. 2016).

### Tissue Collection and Processing

Pups were weighed and sacrificed, while embryos were collected by dissection in ice-cold PBS, excess PBS drained and whole embryos weighed. Brains were then dissected from the skull in ice-cold PBS and cut below the brain stem, immediately drained on paper towels using a slotted spoon and wet weights determined using a fine scale. Brain weights were normalised to body weight and group differences were calculated using unpaired student’s t-test. Brains were post-fixed in 4% PFA at 4°C for 24h, dehydrated and paraffin embedded. Serial coronal sections were cut at 10μm such that each slide contained three consecutive sections.

#### Immunohistochemistry

Sections were re-hydrated using standard protocols and heated in 10mM Sodium Citrate solution (pH6). Endogenous peroxidases were blocked by incubating in 3% H_2_O_2_ and 10% MeOH in PBS for 15mins. Sections were permeabilised in 0.2% Triton X-100 (Sigma Aldrich) in PBS (PBT2) for 5 mins and blocked using 10% heat-inactivated normal goat serum and 2% gelatin in PBT2 for 1 hour. Sections were incubated in 5% GS in PBT2 containing primary antibody (rabbit anti-phosphohistone 3B (Cell Signaling (#9701), 1/100)). overnight at 4°C. After incubation with primary antibody, sections were incubated in biotinylated anti-rabbit immunoglobulin secondary antibody (Dako (#E0432), 1/200) in 5% goat serum in PBT2. Samples were washed in PBS and incubated with Avidin/biotin complex (ABC, Vector) in PBS for 1 hour. Sections were developed using 0.025% DAB and 0.03% H_2_O_2_ in PBS for 10 mins, counterstaining using Ehrlich’s Hematoxylin solution and mounted in DPX (Sigma-Aldrich). Images were acquired on a Nikon 80i microscope equipped with a Nikon 5M pixel Nikon DS digital camera. Images were processed using Adobe Photoshop and Illustrator.

### RNA Extraction and qRT-PCR Analysis

To extract RNA, dissected cortices were lysed in 600μl Trizol (Life Technologies). RNA was purified and DNase-treated using the Direct-zol RNA MiniPrep kit (Zymo Research) according to the manufacturer’s instructions. For qRT-PCR, cDNA was synthesized using 50ng RNA from 4 biological replicates per condition with the Precision nanoScript 2 Reverse Transcription Kit (PrimerDesign Ltd.) according to the manufacturer’s recommendations. qRT-PCRs were performed on a Stratagene Mx3000p (Agilent Technologies) using PrecisionPlus-MX 2x qPCR Mastermix with SYBR green (PrimerDesign Ltd.) and primers against *Chd8* exon 3-4 (FW: CAG AGG AGG AGG GTG AAA AGA AAC, Rev: GAG TTG TCA GAC GAT GTG TTA CGC) or *Chd8* exon 1-2 (FW: TGA AGC CTG CAG TTA CAC TGA CGT, Rev: CTG CGG CTG TGG CTG TGG TT). *Canx* and *Sdha* (E12.5) and *Gapdh* and *Eifa* (P5) were used as endogenous control genes as determined by prior geNorm (Primerdesign, UK) analysis for the respective sample sets. Relative expression levels were calculated using the 2^-ΔΔC_T_^ method.

### RNA Sequencing

RNA was isolated from micro-dissected cortices at E12.5 (both hemispheres) and P5 (one hemisphere) and reverse transcribed (n=3 per experimental group). cDNA was end-repaired, adaptor-ligated and A-tailed. Paired-end sequencing was performed on the Illumina HiSeq 4000 platform. Quality of the raw sequencing data was checked using FastQC version 0.11.2 (Andrews 2010; available at: http://www.bioinformatics.babraham.ac.uk/projects/fastqc) and trimming of adaptor sequences was performed using Trim Galore! version 0.4.1 (Krueger 2012; available at: http://www.bioinformatics.babraham.ac.uk/projects/trim_galore/). Reads were aligned to the mouse genome (GRCm38.p4) using Tophat version 2.1.0 and aligned reads were counted using FeatureCounts version 1.5.0 (Kim et al. 2013; Liao et al. 2014). Differential expression testing was performed using DESeq2 version 1.10.1, as previously described (Love et al. 2014). Gene ontology analysis and functional classification was performed using DAVID with all detected DEGs below a 0.05 FDR (Huang da et al. 2009). Putative regulatory transcription factors were determined with Enrichr using the “ENCODE and ChEA Consensus TFs from ChIP-X” database with all DEGs below a 0.05 FDR (Chen et al. 2013). The R package ggplot2 version 2.1.0 was used to generate volcano plots and DESeq2 normalised read count plots for individual genes. The list of ASD associated genes used for overlap with P5 DEGs was obtained from the SFARI Gene Scoring module (https://gene.sfari.org/autdb/HGHome.do). RNA-seq data have been deposited into GEO, accession number GSE81103.

### Gene Expression Enrichment Analysis

4345 gene expression images corresponding to 4082 unique genes were downloaded from the Allen Institute’s Mouse Brain Atlas coronal expression dataset (Lein et al. 2007). The coronal expression dataset is limited by the fact that it only partially covers the genome and was defined in a biased manner (Ng et al. 2009). However, it offers higher data quality and resolution especially in lateral cortical areas including auditory cortex, compared to the sagittal dataset. This dataset, obtained via in situ hybridization, consisted of 3D spatial expression images aligned to a single reference model and summarized over the whole mouse brain at a 200μm isotropic resolution. Specifically, the gene expression energies, defined by the Allen Institute as the sum of expressing pixel intensities divided by the sum of all pixels in a subdivision, were obtained. Expression data were obtained from adult (P56) male C57B1/6.

Mean gene expression energies were extracted under a set of Allen Institute-defined segmentations, resulting in vectors of expression values for each gene that describe their spatial expression patterns. To account for differences in probe affinities that subsequently affect the total expression levels reported in the images, expression values for each gene were further normalized by dividing by the total expression (summed over regions) for that gene.

Next, we extracted rsfMRI time series data under the aforementioned set of segmentations. This was achieved by aligning, via ANTS (Avants et al. 2009; available at: http://hdl.handle.net/10380/3113), the Allen Institute’s average two-photon microscopy template over which the parcellations were defined to a high resolution (56μm) T2-weighted MR template of the mouse brain, resulting in an atlas defined on MRI data. This atlas was then combined with the MAGeT procedure (Chakravarty et al. 2013) implemented in the PydPiper framework (Friedel et al. 2014) to generate segmentations for each of the T2 weighted images acquired during the course of the rsfMRI experiments. The strongest overconnectivity under these parcellations was observed between the CA2 region of the hippocampus (“CA2”) and the auditory cortex (“AUD”), confirming our previous findings.

We then determined candidate genes that may be associated with the functional overconnectivity phenotype by choosing those that had the highest normalized expression in both regions (“CA2” and “AUD”), compared to all other genes (Supplementary Fig. 4A). Candidates that were in the top 20% of genes for both regions were passed through an enrichment analysis via the GOrilla tool (Eden et al. 2009; http://cbl-gorilla.cs.technion.ac.il/). Another set of candidates were obtained by ranking each gene by the sum of their normalized expression values in both regions (equivalent to the L1 distance); this set also favours genes with high expression in one of the two regions, along with both regions (Supplementary Fig. 4B). For enrichment analyses of ranked lists of genes, GOrilla automatically thresholds the list independently for each GO term so that the optimal enrichment is found (Eden et al. 2009).

To establish specificity of the enriched gene ontology terms, we performed equivalent analyses between 10 randomly chosen regional pairs (DG – FRP, LA – MB, OLF – IG, BLA – P-sat, MY-sat – DG, PAL – GU, P-sat – LZ, PAA – VERM, AON – HEM, MY-mot – MB). The number of returned GO terms was generally low and did not show the enrichment for neuronal development terms seen for the CA2 – AUD pair.

### Statistical Analysis

Data are reported as Mean±SEM and graphs show all individual data points where feasible. Significant p-values are reported in the results section and figure legends provide full details of all relevant statistical parameters including group sizes. Statistical analyses were performed either with SPSS (Version 22, IBM, Armonk, USA) or GraphPad Prism (Version 6, GraphPad Software, La Jolla, California, USA). All analyses were performed blind to genotype.

#### Behaviour

Data were analysed using either a between-subjects ANOVA or a 2-way repeated measures ANOVA, as appropriate. If there was no statistically significant sex difference, data were pooled. When the appropriate ANOVA showed a significant effect for a particular task, student’s t-tests were used as post-hoc analyses, as there were only 2 groups for comparison. Cohort details can be found in the methods, group sizes are stated in the figure legend.

#### Proliferation

Phosphohistone 3B-positive cells lining the ventricular surface of the dorsal cortex were counted and normalised to the length of ventricular surface. These were quantified on both sides of the brain in three consecutive sections and averaged to calculate the number of phosphohistone 3B-positive cells per μm of ventricular surface in the dorsal cortex. Group differences were calculated using unpaired student’s t-test.

#### μCT analysis

Each 3D landmark point was recorded twice for each sample and distances between landmark points normalised to the average of the wildtype controls. Group differences for distances between two specific 3D landmark points were calculated using unpaired student t-test.

#### MRI analyses

Processing of raw data is described in detail in the relevant method sections. For structural MRI, significant differences were determined between groups for the 159 different regions in the brain. Voxelwise comparisons were made between mutants and littermate controls, group differences were calculated using unpaired student’s t-test and multiple comparisons were controlled for using a False Discovery Rate (FDR< 0.15). Exact p-values can be found in Supplementary Table 1.

For rsfMRI studies, group-level differences in connectivity distributions were calculated using 2-tailed student’s t-test (p<0.05, family-wise error cluster-corrected, with cluster defining threshold of t_24_>2.06, p<0.05) and multiple comparisons were controlled for using an FDR<0.05.

#### RNAseq

Processing of raw data and differential expression testing is described in the methods section. Multiple comparisons were controlled for using an FDR<0.05. Exact p values and FDR adjusted p-values for all differentially expressed genes are listed in Supplementary Tables 2 & 3.

## RESULTS

A mouse line with a conditional *Chd8* allele was produced through homologous recombination in C57Bl/6J embryonic stem cells (Supplementary Fig. 1A, B). *Chd8^flox^* mice were crossed with the ubiquitously expressing *βactin-Cre* line (Lewandoski and Martin 1997) to generate *Chd8^+/-^* mice (Supplementary Fig. 1C). Cre-mediated deletion of loxP-flanked (flox) exon 3 results in an early frameshift and termination of translation at amino acid 419, predicted to produce a protein that lacks all functional domains, equivalent to nonsense and frameshift mutations terminating CHD8 at amino acids 62 and 747 in patients (Barnard et al. 2015).

Quantitative RT-PCR (qRT-PCR) on RNA isolated from E12.5 and P5 neocortices using primers spanning the exon 3/4 boundary showed *Chd8* expression reduced by 64% (p=0.006) and 52% (p=0.01), respectively (Supplementary Fig. 1D). CHD8 protein levels were reduced by 51% in *Chd8^+/-^* E12.5 neocortices compared to controls (Supplementary Fig. 1E,F), validating our *Chd8^+/-^* mice as a suitable model for *CHD8* haploinsufficiency. Importantly, we found no evidence for a truncated protein product of 419aa (∼45kDa) that may have resulted from translation of any mutant transcript (Supplementary Fig. 1E). qRT-PCR analysis at E12.5 with primers spanning the exon 1/2 boundary (upstream of the recombination event) revealed reduced *Chd8* expression of 52% (Supplementary Fig. 1G), indicating that the mutant transcript is most likely subject to nonsense-mediated decay.

### *Chd8* heterozygous mice have specific craniofacial and structural brain phenotypes

Humans with truncating mutations in a single *CHD8* allele often present with macrocephaly (64%) and distinct craniofacial phenotypes (89%), which include hypertelorism (wide-set eyes, 67%) (Bernier et al. 2014; Stessman et al. 2017). We characterised the cranioskeleton of *Chd8^+/-^* mice by μCT to ask whether these phenotypes were also present in *Chd8^+/-^* mice (Fig. 1A–D). The interorbital distance (landmarks 8-9, Fig. 1C,D) was significantly wider in *Chd8^+/-^* mice compared to controls, indicative of a hyperteloric phenotype (*p=0.0273; Fig. 1C,D,F). In addition, the anterior-posterior length of the interparietal bone (landmarks 4-5) is increased in *Chd8^+/-^* animals (**p=0.0025; Fig. 1A,B,E), suggestive of more wide-spread craniofacial anomalies associated with *CHD8* haploinsufficiency.

**Figure 1.**
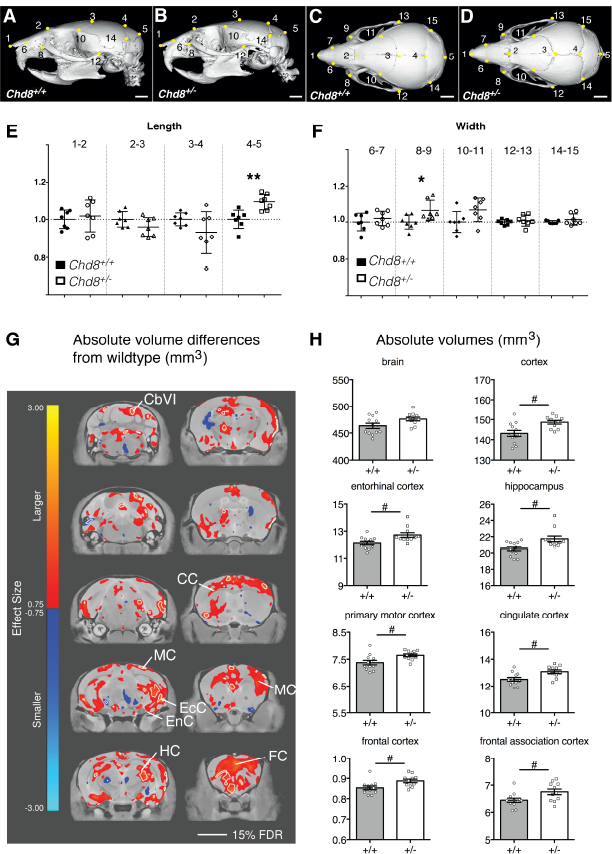
Hypertelorism and mild megalencephaly in *Chd8^+A^* mice. A-D) Representative lateral (A,B) and dorsal (C,D) μCT views of 3D reconstructed skulls from mice with the indicated genotypes. Landmarks from 1 to 15 are indicated by yellow dots. Scale bars = 2mm. E,F) Graphs for measurements between indicated landmarks, normalised to average measurements from corresponding wildtype littermates. Mean±SEM; landmarks 4-5: p=0.0025, t=3.797; landmarks 8-9: p=0.0273, t=2.512; df=12, student’s t-test, n=7 per genotype. G) High-resolution 7T structural MRI coronal images of *Chd8^+/-^* brains from posterior (top left) to anterior (bottom right) are shown. Absolute volumetric differences in size, relative to wildtype controls are coloured according to the scale on the left. Effect size is measured in units of standard deviation. Some regions with enlarged volumes are labelled as follows: CbVI – cerebellar lobule VI, MC – motor cortex, EcC – ectorhinal cortex, EnC – entorhinal cortex, HC – hippocampus, CC – Cingulate cortex, FC – frontal association cortex. H) Absolute volumes (mm^3^) are plotted for whole brain, neocortex and several other brain regions for the different genotypes as indicated. #FDR<0.15, student’s t-tests: brain: p=0.0484, t=2.096; cortex: p=0.0055, t=3.093; entorhinal cortex: p=0.011, t=2.788; hippocampus: p=0.0091, t=2.873 primary motor cortex: p=0.0126, t=2.727; cingulate cortex: p=0.0074, t=2.965; frontal cortex: p=0.0154, t=2.639; frontal association cortex: p=0.0238, t=2.438; df=21, *Chd8^+/-^:* n=11, *Chd8^+/+^:* n=12. Individual volumes and volume differences for all brain regions are listed in Supplementary Table 1.

To examine whether structural brain abnormalities were present in *Chd8^+/-^* mice, their brains were compared to *Chd8^+/+^* littermates by high resolution MRI (Fig 1G). Total brain volume was increased by 2.7% in *Chd8^+/-^* mice (476mm^3^ vs. 463mm^3^, p=0.048, FDR=15%, Fig. 1H). Accordingly, several brain regions, including cortical areas, hippocampus and parts of the cerebellum showed volumetric increases (Fig. 1G, H, Supplementary Table 1). Structural alterations of these brain areas have been implicated in autism (Blatt 2012; Donovan and Basson 2017; Ecker 2016) providing potential neural substrates for the autism phenotype associated with *CHD8* haploinsufficiency in humans.

### *Chd8^+/-^* mice show abnormal activity levels and differences in social interaction

We next assessed *Chd8^+/-^* mice in a number of behavioural tests to ask whether they exhibited any signs of socio-communicative deficits, repetitive behaviours or cognitive inflexibility, representing core ASD-like behaviours in humans.

*Chd8* heterozygous pups displayed signs of delayed motor development in the first two weeks after birth. *Chd8^+/-^* pups took slightly longer than wildtype littermates to develop an effective righting reflex over time (*p=0.014; Fig. 2A). Correspondingly, *Chd8^+/-^* pups spent more time engaged in unsuccessful attempts to turn over on their stomachs as measured during the spontaneous motor behaviour observations (P6: *p=0.0312, P8: *p=0.0354; Fig. 2B). Once they were able to move around the cage, mutant pups spent on average more time in locomotion than wildtype littermates suggestive of hyperactivity (**p=0.009; Fig. 2C).

**Figure 2.**
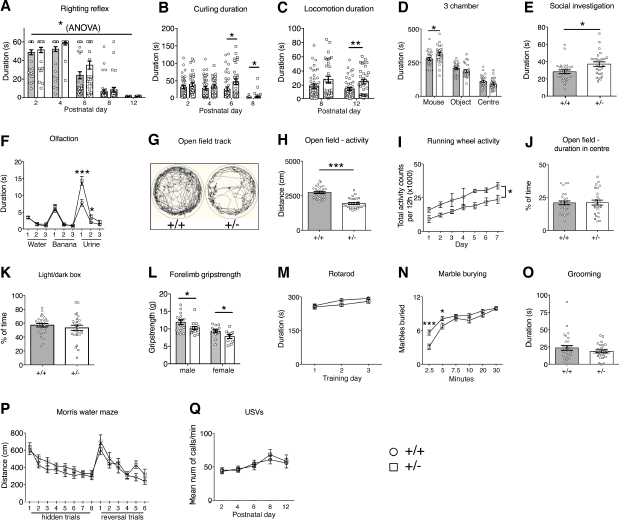
Complex behavioural abnormalities in *Chd8* heterozygous mice. A-Q) Behavioural assessments of a cohort of adult *Chd8^+/-^'* (+/-, n=16 (male), n=14 (female)) and *Chd8^+/+^* (+/+, n=15 (male), n=14 (female)) and of pup *Chd8^+/-^'* (+/-, n=11 (male), n=20 (female)) and *Chd8^+/+^* (+/+, n=22 (male), n=20 (female)) animals. A) The development of the righting reflex in pups at the indicated postnatal days. Pups failing to right by the end of the 60s test period were given a score of 60s. Note the significant delay in the acquisition of the full righting reflex response in *Chd8+'* animals compared to littermate controls. Mean ±1SEM; *p=0.014 (one-way repeated-measures ANOVA: f(1,72)=6.36 (between-subjects effect)). B) The duration, in seconds, pups spend rolling on their back (curling) as recorded during the analysis of spontaneous movements during USV recordings. Note that *Chd8^+/-^* mice spent significantly more time curling at P6 and P8 compared to littermate controls. Mean ± SEM; P6 *p=0.0312, P8 *p=0.0354 (one-way repeated-measures ANOVA: f(1,72)=12.64 p=0.001 (between-subjects effect), with student’s t-test as post-hoc analysis (p6: df=72, t=2.197, P8: df=72, t=2.145). C) The duration, in seconds, pups spend in locomotion as recorded during the analysis of spontaneous movements during USV recordings. At P12 *Chd8^+/-^* animals spent significantly more time in locomotion as compared to littermate controls. Mean ±1SEM, **p=0.009 (oneway repeated measures ANOVA: (f(1,72)=7.33 p=0.008 (between-subjects effect), with student’s t-test as post-hoc analysis df=72, t=2.687). D) The duration, in seconds, spent in each chamber of the three-chamber sociability test. All mice spent a significantly higher proportion of time in the chamber with the age and sex matched stranger con-specific mouse compared to the other chambers. Mean±SEM; *p=0.029 (between-subjects ANOVA: f(1,53)=5.031). E) Duration, in seconds, of social investigation over a three-minute period. Social investigation was defined as the combined total duration of head, body and anogenital sniffing of a conspecific mouse. Mean ±SEM; *p=0.015 (between-subjects ANOVA f(1,52)= 6.307). F) Graph demonstrating the performance in the olfactory habituation/dishabituation test. Mean±SEM; *p=0.03, **p=0.0002 (repeated-measures ANOVA: f(2.85,145.23)=9.24, p=0.00002, with student’s t-test as post-hoc analysis **df=53,t=4.04, *df=53,t=2.23). G) Representative ethovision tracks of a *Chd8^+/-^* (+/-) and *Chd8^+/+^* (+/+) animal plotting their movements during the 10-minute open field task. H) The total distance travelled in the outer part of the open field arena over a 10-minute time period. Mean±SEM; ***p=2×10^−9^ (between-subjects ANOVA: f(1,53)=52.72). I) The total activity counts per 12h period on running wheels in the homecage during 7 days of dark-phase recording. Mean±SEM; *p=0.019 (repeated-measures ANOVA: f(1,7)=9.12, between-subjects effect). J) The percentage of time spent in the centre of the open field arena during the 10-minute test. Mean±SEM (between-subjects ANOVA: f(1,53)=0.007, p=0.93). K) The percentage of time spent in the light chamber during the 5minute light/dark test. Mean±SEM (between-subjects ANOVA: f(1,51)=0.824, p=0.368). L) The average of 3 measurements of forelimb grip strength on a Linton Grip Strength meter. Mean ± SEM, males: *p=0.045 (between-subjects ANOVA: f(1,29)=4.371) females: *p=0.042 (between-subjects ANOVA: f(1,22)=4.677). M) The mean latency of mice to fall from the rotarod. Mean±SEM (repeated-measures ANOVA: f(1.644,102)=0.620, p=0.540). N) The average number of marbles buried, out of a maximum of 12, within a 30-minute time period. Mean±SEM; *p=0.04, ***p=0.0004, (repeated-measures ANOVA: f(3.66,265)=4.70 p=0.002, with student’s t-test as post-hoc analysis * df=53,t=2.12, ***df=53, t=3.79). O) The duration, in seconds, mice spent self-grooming during the 10-minute self-grooming test. Mean±SEM (between-subjects ANOVA: f(1,51)=1.21, p=0.28). P) Graph plotting the average distance swum for 4 trials daily over 8 consecutive training days to find the hidden platform (hidden trials), followed by 6 training days where the location of the platform was reversed (reversal trials). Mean±SEM (repeated-measures ANOVA: f(8.761,714)=1.064, p=0.388). Q) The mean number of ultrasonic vocalisations per minute on indicated postnatal days. Mean±SEM (repeated-measures ANOVA: f(1,72)=0.76, p=0.39).

In the three-chamber sociability test, adult *Chd8^+/-^* mice spent significantly more time in the chamber with the novel age and sex-matched conspecific mouse than in the other chambers, indicative of normal sociability (Fig. 2D). Interestingly, rather than displaying sociability deficits, mutant mice spent slightly, but significantly more time in the chamber containing the mouse, compared to controls (*p=0.029; Fig. 2D). *Chd8^+/-^* mice also spent more time investigating conspecific mice in a reciprocal social interaction test (*p=0.015; Fig. 2E). A quantitative olfactory habituation/dishabituation test revealed an increased interest in an odour with social significance (urine) in *Chd8^+/-^* mice compared to controls (*p=0.03, ***p=0.0002; Fig. 2F). No difference in the time spent investigating a non-social (banana) odour was observed, implying an increased interest specifically in social cues and an otherwise normal capacity for odour discrimination (Fig. 2F).

Examination of these animals in the open field arena revealed a marked hypo-activity in *Chd8^+/-^* mice (***p=10^−9^; Fig. 2G,H). The hypo-active phenotype was also observed in mutant mice in their homecage environment by measuring activity on a running wheel over a one-week period (*p=0.019; Fig. 2I). The open field test did not show any evidence of anxiety in these mice, i.e. an increased reluctance to enter the inner, most exposed area of an open field arena (Fig. 2J). This was confirmed in the light/dark box test that showed no difference between wildtype and mutant mice (Fig. 2K). Forelimb grip strength was slightly but significantly reduced in mutant mice (*p=0.045 (males) *p=0.042 (females); Fig 2L) but *Chd8^+/-^* mice showed normal motor abilities on the revolving rotarod, indicating that a reduced capacity to perform motor tasks was unlikely to be the cause of the hypo-active phenotype (Fig. 2M). No evidence of repetitive behaviours was observed by assessing marble burying and self-grooming behaviours (Fig. 2N,O). In fact, mutants showed slightly delayed marble burying behaviour, most likely due to their general hypoactivity (*p=0.04, ***p=0.0004; Fig. 2N).

Spatial learning abilities and cognitive flexibility were assessed in the hippocampus dependent Morris water maze test. *Chd8^+/-^* mice performed normally in the learning part of this test (Fig. 2P). In a reversal paradigm, these mice were also indistinguishable from wildtype littermates, implying normal cognitive, spatial learning abilities and flexibility (Fig. 2P). Finally, no differences in the number of ultrasonic vocalisations (USVs) of pups separated from the nest were recorded, indicating no obvious communication deficits (Fig. 2Q).

As male mice from this adult behavioural cohort were used for structural MRI analyses (Fig. 1G, H) we were able to correlate their brain volume with specific behaviours. We observed significant inverse correlations between activity in the open field and hippocampal volume (*p=0.012; Supplementary Fig. 3A), as well as cortical volume (*p=0.036; Supplementary Fig. 3B). Overall brain volume and hypoactivity showed a weaker correlation (p=0.092; Supplementary Fig. 3C) hinting at a degree of specificity for cortical and hippocampal regions.

In summary, *Chd8^+/-^* mice displayed no socio-communicative deficits, but rather exhibited a heightened interest in social cues. No evidence for perseverative and repetitive behaviours were observed. *Chd8^+/-^* pups showed evidence for hyperactivity and delayed motor development while adult *Chd8^+/-^* mice exhibited a hypo-active phenotype, which was significantly correlated with overgrowth in cortical and hippocampal regions.

### *Chd8* haploinsufficiency causes general growth delay but postnatal brain overgrowth

To determine whether the brain overgrowth phenotype was already present at early postnatal stages when developmental delay was evident, we measured body and brain weights from birth. *Chd8^+/-^* pups showed significant growth retardation from postnatal day 4 onwards and into early adulthood (Fig. 3A). Brain and body weight were well correlated in both wildtype and heterozygous mice at P35 (r^2^=0.25, p=0.0004 and r^2^=0.28, p=0.005, respectively), with *Chd8* mutants displaying higher brain weights compared to their wildtype littermate controls with equivalent body weight (Fig. 3B). A group-wise comparison confirmed the significant increase in normalised brain weight in *Chd8^+/-^* mice compared to wildtype littermates (20.4% increase, ***p<0.0001; Fig. 3C). At P7, normalised brain weights were already significantly larger in *Chd8^+/-^* pups compared to wildtype littermate controls (9.3%, ***p=0.0009) with more subtle differences between the groups observed at P0 (6.7%, *p=0.01).

**Figure 3.**
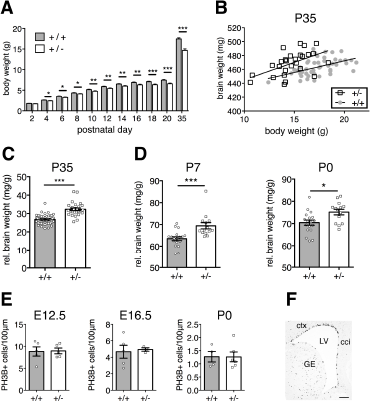
*Chd8* mutants display postnatal brain overgrowth. A) Body weights of mice between P2 and P35. Repeated-measures ANOVA with student’s t test as post hoc analysis. ANOVA: f(2.303, 158.940)=12.313, p=0.000003; student’s t-tests: P4: t=2.498, *p=0.0148; P6: t=2.385, *p=0.0197; P8: t=1.916, *p=0.0593; P10: t= 2.808, **p=0.0064; P12: t= 2.803, **p=0.0065; P14: t=2.018, **p=0.0035; P16: t=3.353, **p=0.0013; P18: t=4.082, ***p=0.0001; P20: t=4.269, ***p<0.0001; P35: t=6.334, ***p<0.0001; df=71, +/+, n=46; +/-, n=27. B) Individual wet brain weights plotted against individual body weights for *Chd8^+^*^-^ (+/-), mice and their littermate controls (+/+) at postnatal day (P)35. Note that *Chd8^+^'∼* mice have larger brain weights than littermate controls of equivalent body weight. Linear regression analysis: r^2^=0.25, p=0.0004 (+/+), r^2^=0.28, p=0.005 (+/-). C) Wet brain weights normalised to body weight P35. *Chd8^+-^* (+/-) show significantly increased normalised brain weights compared to their littermate controls (+/+). Mean ± SEM; ***p<0.0001, t=7.455 (student’s t-test). +/+, n=46; +/-, n=27. D) Wet brain weights of pups at P7 and P0 normalised to body weight. *Chd8^+^*^-^ (+/-) pups show significantly larger normalised brain weights than their littermate controls (+/+) at P7 and P0. Mean ± SEM; *p=0.01, t-2.746; ***p=0.0009, t=3.681 (student’s t-test). P7: +/+, n=18; +/-, n=14; +/+; P0: +/+, n=18; +/-, n=14; +/+. E) Quantification of phospho-histone H3 (PH3B) positive cells in the ventricular zone at E12.5, E16.5 and P0. Cell counts were normalized to ventricular surface length. Mean ± SEM; student’s t-test. E12.5: +/+, n=5; +/-, n=5. E16.5: +/+, n=5; +/-, n=3. P0: +/+, n=4; +/-, n=6. F) Example of PH3B immunostaining in an E16.5 coronal brain section. Scale bar=100μm; LV: lateral ventricle, ctx: cortex, cci: cingulate cortex, GE: ganglionic eminence.

Together, these analyses suggested that subtle, but cumulative differences in brain growth over several days may be responsible for small increases in brain size. Indeed, we did not detect any significant differences in cortical ventricular zone (VZ) proliferation as measured by phospho-histone H3 immunostaining at E12.5, E16.5 or P0 (Fig. 3E,F). However, subtle increases in progenitor proliferation in the VZ cannot be completely ruled out.

### CHD8 controls the expression of ASD-associated axon guidance genes in the early postnatal neocortex

To gain insights into the transcriptional programmes that may underlie the subtle brain overgrowth and abnormal behaviours observed in *Chd8^+/-^* mice, we performed RNA-seq analysis on dissected neocortical tissue at two stages: 1) At E12.5, when *Chd8* expression peaks (Durak et al. 2016) and neural progenitor cells predominate, and 2) At P5, when many developmental processes with relevance for ASD aetiology, such as axon growth and guidance and synaptogenesis, are under way.

Surprisingly, only 5 genes, including *Chd8,* showed significant (FDR<0.05) differential expression in *Chd8^+/-^* embryos at E12.5 in this experiment (Fig. 4A, Supplementary Table 2). By contrast, 649 differentially expressed genes (FDR<0.05) were identified in the P5 neocortex, with over two thirds of these genes downregulated (Fig. 4B, Supplementary Table 3).

**Figure 4.**
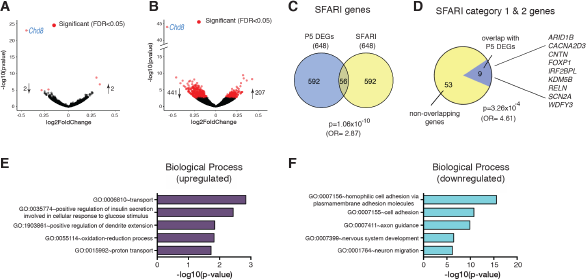
Gene expression changes in *Chd8*-deficient neocortices. A) Volcano plot of RNA-seq data from embryonic (E12.5) *Chd8^+I^* neocortex. Each point represents an individual gene and all genes differentially expressed in *C,hd8^+/^'* samples with an FDR of 0.05 are highlighted in red. All differentially regulated genes are listed in Supplementary Table 2. B) Volcano plot indicating differentially expressed genes (DEGs) detected by RNA-seq in P5 *C'hdH* "neocortex. All differentially regulated genes are listed in Supplementary Table 3. C) Venn diagram showing extent of overlap between P5 DEGs and ASD associated genes (categories 1-5 & S) in the SFARI gene database. Enrichment was calculated using Fisher’s exact test for count data. SFARI genes that overlap with P5 DEGs are listed by category in Supplementary Table 4. D) Pie chart showing the proportion of high confidence ASD candidate genes (categories 1-2) that are found in the P5 DEG set. Enrichment was calculated using Fisher’s exact test for count data. E, F) Results of gene set enrichment analysis using the DAVID knowledgebase on the P5 DEG set (FDR<0.05). The five most significant Gene Ontology terms in the Biological Processes category are shown for up-regulated DEGs (E) and downregulated DEGs (F), respectively. The five most significant Gene Ontology terms in the molecular function and pathways categories and the four most over-represented transcription factors identified by Enrichr analysis are shown in Supplementary Fig.2. A comprehensive list of all significant Gene Ontology terms in the biological processes, molecular functions and pathways categories is given in Supplementary Tables 5 to 10.

Comparing all differentially expressed genes (DEGs) from the P5 dataset with the SFARI autism gene list identified 56 shared genes, representing a highly significant enrichment of ASD-associated genes in the differentially expressed gene set (p=1.06×10^−10^ (OR=2.87); Fig. 4C, Supplementary Table 4). Almost all (53/56 = 95%) of these ASD-associated genes were down-regulated (Supplementary Table 4). We also overlapped our gene set with high confidence (SFARI categories 1&2) ASD candidates (p=3.26×10^−4^ (OR=4.61); Fig. 4D). Nine genes, representing 16% of all SFARI category 1 & 2 genes, were present in our differentially expressed gene set at P5. All of these high confidence ASD candidate genes were down-regulated (Supplementary Table 4).

Amongst the upregulated gene set, the most significant KEGG pathways, molecular functions and biological processes were related to the ribosome and oxidative phosphorylation, whereas the downregulated gene set included categories related to cell adhesion, axonal guidance and calcium signaling pathways (Fig. 4E, F, Supplementary Fig. 2A, Supplementary Tables 5-10). Identification of potential regulatory transcription factors was performed using Enrichr, which found over-representation of Suz12 targets in the down regulated gene set (Supplementary Fig. 2B). Suz12 is a component of the Polycomb repressor complex 2 (PCR2) and is required for both histone methyl transferase and gene silencing activities of PRC2 (Cao and Zhang 2004). The observation that Suz12 targets are overrepresented in the down-regulated gene set offers a potential mechanistic explanation for the down-regulation of some of the identified genes. None of the genes that code for components of PRC2, including *Suz12,* are differentially expressed at P5, suggesting that the enrichment seen was unlikely to be due to direct transcriptional dysregulation of polycomb gene expression at this stage of development.

### *Chd8^+/-^* mice exhibit over-connectivity in cortical and hippocampal networks

The significant enrichment of cell adhesion and axonal guidance genes in the down-regulated gene set at P5 led us to hypothesise that long-range connectivity might be disrupted in *Chd8* heterozygous neocortices. To test this hypothesis, we performed rsfMRI to probe functional brain connectivity in mature brain networks. Synchronous fluctuations in blood-oxygen-level dependent (BOLD) signals in different brain regions are used as an indication of them being functionally connected. A regionally unbiased analysis for long-range connectivity changes revealed hotspots for increased connectivity in *Chd8^+/-^* mice compared to wildtype littermate controls, which included the entorhinal, retrosplenial, auditory cortical and posterior hippocampal areas (t-test, p<0.05 FEW cluster-corrected, with cluster-defining threshold t_24_>2.06, p<0.05; orange areas in Fig. 5A). This analysis suggested that hyper-connected areas were predominantly located on the left side of the brain. A re-analysis of these results without the use of cluster correction revealed the presence of foci with increased connectivity also on the right side, mirroring the effects observed on the left (dark red areas in Fig. 5A). Inter-hemispheric mapping of rsfMRI connectivity strength in previously characterized rsfMRI network systems of the mouse brain (Sforazzini et al. 2014), revealed increased cortical connectivity in auditory regions (p<0.05, student’s t-test, uncorrected), although the effect did not survive false discovery rate correction (q=0.05) for multiple comparison across the rsfMRI networks probed. We next used a seed-based approach to specifically probe regions with altered connectivity to these hotspots to reveal the brain networks affected. Most strikingly, this revealed a reciprocal increase in connectivity between ventral hippocampus and auditory cortical regions in *Chd8* mutant mice (t-test, p<0.05 FEW cluster-corrected, with cluster-defining threshold t_24_>2.06, p<0.05; Fig. 5B,C). Seed placement in the auditory cortex revealed increased connectivity of this region with both cingulate and entorhinal cortices (Fig. 5B), whereas a hippocampal seed uncovered strengthened long-range connectivity with somatosensory and visual cortices (Fig 5C).

**Figure 5.**
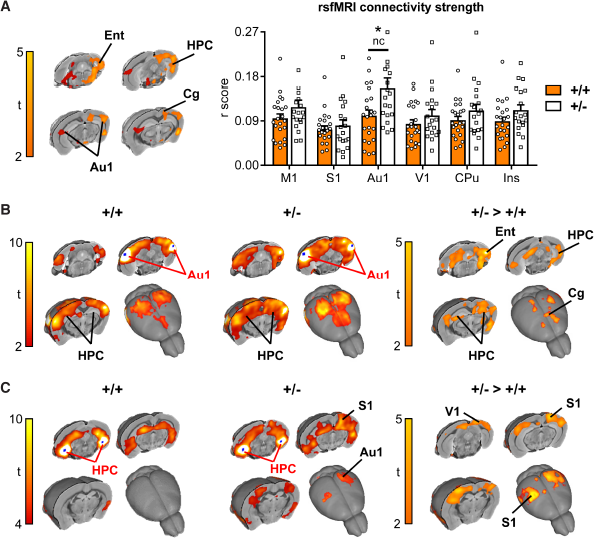
Resting-state functional MRI reveals increased parieto-hippocampal connectivity in *Chd8+'* mice. A) Global (long range) connectivity mapping revealed bilateral foci of increased connectivity in posterior cortical and hippocampal regions in *Chd8+^/^* mice with respect to control littermates (*Chd8^+/-^:* n=19, *Chd8^+/-^* +: n=23). The effect is shown at a threshold t24>2.06, P<0.05 (Fig 5A). Orange areas indicate regional rsfMRI differences surviving cluster correction (p=0.05). The bar plot on the right displays global connectivity strength quantification (r *score)* in bilateral areas of interest. *<p0.05, nc=not corrected for multiple comparisons. Ent – entorhinal cortex, HPC – hippocampus, Au1 – auditory cortex, Rs – retrosplenial cortex, M1 – motor cortex, S1 – somatosensory cortex, V1 – visual cortex, CPu – caudate putamen. Ins – insular cortex. B) Seed based connectivity mapping obtained using a bilateral auditory seed (Au1) covering foci of increased global rsfMRI connectivity depicted in A. A robust increase in rsfMRI connectivity was observed in hippocampal (HPC), entorhinal (Ent) and cingulate (Cg) regions of *Chd8^+/-^* mice (t<2, pc=0.01). C) Seed based connectivity mapping obtained using a bilateral ventro-hippocampal seed (HPC) covering bilateral foci of increased global rsfMRI connectivity in A. A significant increase in rsfMRI connectivity was observed in peri-hippocampal and auditory/parietal (S1 and V1) regions in *Chd8^+/-^* mice (t<2, pc=0.01).

Previous studies suggested that neurodevelopmental genes show correlated expression patterns in brain areas that are connected (French and Pavlidis 2011). To test if the same applies to our model we harnessed gene expression data contained within the Allen brain atlas to identify genes that are highly expressed in hippocampal and auditory areas. Using expression data from the Allen brain atlas, we determined genes with high relative expression in both the hippocampal CA2 region and auditory areas (Supplementary Fig. 4A,B; Supplementary Table 11; see methods for details). Enriched genes preferentially clustered into GO term categories relating to synaptic development and function, axonal structure and neuron projections (Supplementary Table 11). To determine if any of these highly expressed genes are likely contributing to a developing process linked to the functional connectivity phenotype, we compared them with the DEG set from our P5 RNAseq experiment. This analysis revealed several axon guidance and cell adhesion genes that are preferentially expressed in CA2 and auditory areas at adult stages and whose expression is dysregulated at P5 (e.g. *Cdh2, Cdh11, Cdk5r1, Epha4, Fat3, Nrcam, Robol1;* Supplementary Table 11). Taken together, two independent experimental approaches at different time points identified specific axon guidance and cell adhesion genes, further strengthening our hypothesis that dysregulation of these genes may be important for the functional connectivity phenotype and provides a solid platform for future detailed analyses.

We conclude that early postnatal gene expression changes indeed prefigure abnormal functional connectivity in *Chd8* heterozygous mice. Our findings suggest that abnormalities in specific cortical-hippocampal circuits involved in sensory processing may underlie some of the unique anomalous behaviours observed in *Chd8^+/-^* mice, and by extension, the neuropsychiatric symptoms in patients with *CHD8* mutations.

## DISCUSSION

Here we identify a crucial developmental role for *CHD8* in regulating axon guidance gene expression in the early postnatal period and, for the first time, associate *CHD8* haploinsufficiency with functional over-connectivity of specific brain areas. A recent rsfMRI study involving over 150 male probands with an ASD diagnosis and nearly 200 typically developing individuals described over-connectivity between sensory cortices and subcortical structures as a central feature in ASD (Cerliani et al. 2015). It will be very important to determine whether these specific functional connectivity abnormalities are present in patients with *CHD8* mutations.

While we cannot at this stage establish a direct causal relationship between transcriptional, connectivity and behavioural phenotypes, our data suggest that functionally altered connectivity of sensory cortical areas in *Chd8* mutant mice underpins behavioural phenotypes. In concordance with previous studies (Gompers et al. 2017; Katayama et al. 2016), our data suggest that moderate expression changes of many genes, rather than severe disruption of few genes, cooperate to give rise to phenotypic changes in *Chd8^+/-^* mice. A total of 21 axon guidance genes are downregulated in the early postnatal period in our *Chd8^+/-^* mice (Supplementary Table 6). Therefore, experimental validation of a causal link will be challenging as multiple axon guidance pathways may contribute to the functional connectivity and behavioural phenotypes.

### *Chd8^+/-^* mice as a model for human *CHD8* haploinsufficiency syndrome

CHD8 is one of the highest confidence ASD-associated genes to emerge from recent exome sequencing studies (Bernier et al. 2014; Iossifov et al. 2014; Neale et al. 2012; O’Roak et al. 2012a; Talkowski et al. 2012). We therefore expected *Chd8^+/-^* mice to present with robust, autism-associated behaviours. *Chd8^+/-^* mice displayed delayed motor development and distinctive behavioural anomalies that featured a heightened interest in social cues, but surprisingly did not include repetitive and perseverative behaviours or communication deficits.

In agreement with other published studies (Gompers et al. 2017; Katayama et al. 2016; Platt et al. 2017) we did not observe repetitive behaviours in *Chd8^+/-^* mice. While Katayama et al. reported increased persistence following directional reversal in the T-maze forced alteration test, suggestive of perseverative behaviours, we did not find such evidence in the Morris water maze test. This may be due to the higher complexity of decision making in the Morris water maze compared to the binary choice required by the T-maze. In addition, *Chd8^+/-^* mice consistently did not show any evidence for perseverative behaviours in the marble burying test (Fig. 2N; Gompers et al. 2017; Platt et al. 2017), suggesting that any perseverative behaviours in *Chd8* mutants may be subtle or task specific. *Chd8^+/-^* mice show an apparent heightened interest in social cues, indicating that altering the *Chd8* gene dosage during development can impact socially motivated behaviours (Fig. 2E). An increased duration of contacts in the social investigation test was also seen in two other published behavioural analyses of *Chd8* heterozygous mouse models (Katayama et al. 2016; Platt et al. 2017). Katayama et al. additionally described a reduced duration of active social contacts in *Chd8* mutants, although all test groups showed evidence for very high levels of anxiety, a known behavioural confound. Katayama et al. and Platt et al. further reported normal sociability but minor deficits in social novelty in the three-chamber social approach task in *Chd8^+/-^* mice (Katayama et al. 2016; Platt et al. 2017).

Despite not observing typical ASD-like behaviours, we did detect a delay in early motor development in *Chd8^+/-^* mice (Fig. 2A,B). There is a growing body of evidence suggesting that delayed motor milestones in toddlers predate and predict the emergence and severity of language deficits in later life (Bedford et al. 2016; Chinello et al. 2016). Of note, the only available longitudinal case reports in the literature also describe early motor delay in both patients with *CHD8* haploinsufficiency (Merner et al. 2016; Stolerman et al. 2016).

A key characteristic of autism is restricted behaviours or interests, which often manifest as hyper or hypoactivity to sensory input or unusual interest in sensory stimuli, for example excessive smelling or touching of objects (Constantino and Charman 2016). One may speculate that the excessive smelling of social cues and the increased duration of social contacts observed in our *Chd8^+/-^* mice may be indicative of behavioural abnormalities in these domains.

### Dysregulation of the cortical transcriptome in *Chd8* heterozygous mice

Gene expression analysis showed little evidence for transcriptional dysregulation at mid embryonic stages, but revealed disruption of key developmental processes involved in establishing brain connectivity in the early postnatal neocortex. These data are in agreement with a recent study where Gompers and colleagues only found a handful of genes differentially expressed in bulk forebrain at E12.5 and E14.5, while detecting subtle changes perinatally (E17.5: 89 DEGs, P0: 35 DEGs; FDR<0.05) and more pronounced dysregulation at adult stages (295 DEGs; FDR<0.05) (Gompers et al. 2017).

Many of the transcripts that were dysregulated in the early postnatal period are themselves ASD-associated genes and were predominantly down-regulated. Our gene expression studies therefore provided strong evidence that a variety of genes, pathways and developmental processes implicated in ASD might be dysregulated by *Chd8* haploinsufficiency.

An expanding number of ASD risk genes have roles in axon guidance, synapse development and plasticity (Bourgeron 2015). We detected significant enrichment of genes in these functional categories in our down-regulated gene set, including the major Slit protein receptors *Robol1* and *Robo2, EhpA4* and 5, and cell adhesion molecules such as *L1CAM* and *Cdh2, 5, 8* and *11* (Supplementary Tables 3 & 6). Similarly, Gompers et al. found enrichment for axon growth and guidance factors amongst down-regulated genes in their M3 module (Gompers et al. 2017). Moreover, Sugathan and colleagues showed enrichment for genes associated with the GO terms ‘cell adhesion’, ‘axon guidance’ and ‘neuron differentiation’ amongst down-regulated genes in *CHD8-* deficient human iPSC-derived neural progenitors. This suggests that these important developmental gene sets are regulated by CHD8 in both mouse and human cells (Sugathan et al. 2014).

In sum, our data identify early postnatal development as a key stage at which transcriptional changes caused by *Chd8* heterozygosity may precipitate ASD-related phenotypes. They further indicate that *Chd8* heterozygosity defines a transcriptional programme characterised by diminished expression of key neurodevelopmental regulators that are predicted to affect cellular functions essential for the appropriate wiring of the brain.

### Increased functional connectivity in sensory networks

Significantly, functional connectivity was altered in the adult brain of *Chd8^+/-^* mice. Our rsfMRI analysis found evidence for over-connectivity between sensory regions in the neocortex and limbic cortical regions. Most notably, the auditory cortex showed a global increase in functional connectivity that involved connections to other cortical areas and reciprocal strengthening of connectivity to the ventral hippocampus. It seems likely that altered connectivity is the consequence of some of the disrupted brain wiring pathways uncovered by our RNA-seq experiments. Encouragingly, the expression of several axon guidance and cell adhesion genes, which are dysregulated at P5, is enriched in hippocampus and auditory cortex (Supplementary Table 11); nevertheless, this hypothesis will require further in–depth scrutiny. More importantly, it will be critical to investigate whether these connectivity changes are pertinent to any of the behavioural anomalies in *Chd8* heterozygous mice or the ASD phenotype in patients with *CHD8* haploinsufficiency. The over-connectivity in networks involving the auditory cortex and the hippocampus is intriguing. Auditory processing deficits in ASD are well documented and range from a lack of lateralisation to a general delay in network maturation (Bruneau et al. 1992; Edgar et al. 2015), although the functional behavioural consequences of these deficits are not clear. Furthermore, over responsivity to sensory stimuli is frequently observed in ASD patients, can affect all sensory modalities and appears to be positively correlated with the severity of autistic traits (reviewed in Sinclair et al. 2017; Tavassoli et al. 2014). Although a definitive causal relationship is difficult to establish, it has been hypothesised that sensory over-responsivity may trigger compensatory and avoidance behaviours that promote the emergence of core behavioural autism traits (Marco et al. 2011). In support, tactile hypersensitivity during critical developmental periods has been shown to underlie anxiety and social deficits in a number of genetic ASD mouse models (Orefice et al. 2016). Whether this would be equally the case for other sensory modalities and a general mechanistic feature in the behavioural aetiology of ASD remains an open question.

## ACKNOWLEDGEMENTS

We thank John Whittingham, Alex Donovan and BSU staff for technical assistance and Chris Healy for μCT scans. We acknowledge the High-Throughput Genomics Group at the Wellcome Trust Centre for Human Genetics (funded by Wellcome Trust grant reference 090532/Z/09/Z) for the generation of the RNA sequencing data and Drs. Brian Nieman and Leigh Spencer Noakes for their MRI sequences. We also thank the Allen Institute for Brain Science for providing gene expression data used in this study (© 2004 Allen Institute for Brain Science. Allen Mouse Brain Atlas. Available from: http://mouse.brain-map.org).

---

**Supplementary Figure 1.**
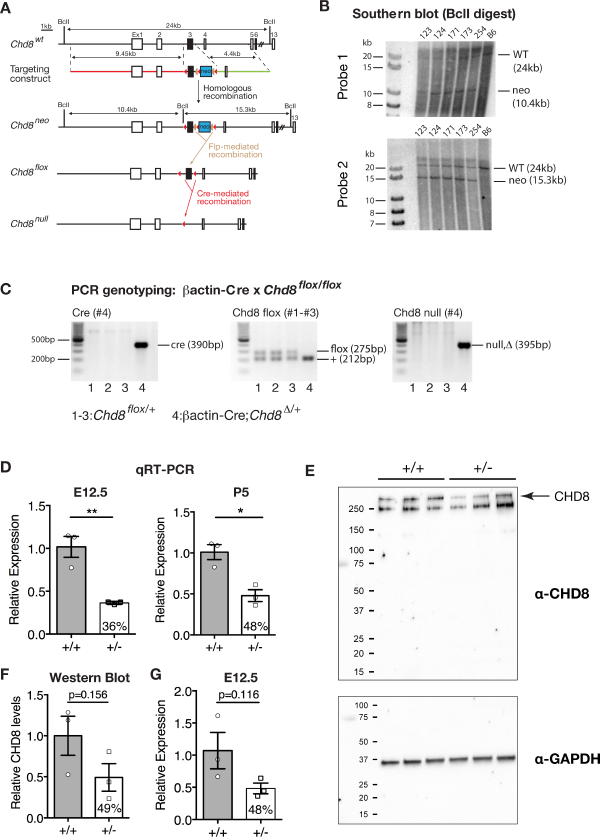
Construction and validation of the *Chd8* conditional and null alleles. A) Diagrammatic representations of the wildtype (wt) *Chd8* locus *(Chd8^wt^),* targeting construct, targeted (neo), conditional (flox) and null alleles. Approximate genomic distances are indicated in kilobases (kb), exons are denoted by boxes labelled Ex1 to 13, and Southern blot probes (P1,P2) and PCR primers (#1 #4) are indicated. The 5’ long homology arm is shown in red and the 3’ short homology arm in green. The neomycin resistance cassette (neo) is shown as a blue box, the floxed exon 3 by a black box, loxP sites by red triangles and frt sites by tan semi-ovals. BclI restriction enzyme sites are labelled. B) Southern blot of genomic DNA digested with BclI from embryonic stem (ES) cell clones (123, 124 etc.) and wildtype C57BL/6J (B6) cells, hybridised with P1 are shown, with molecular weight markers in the left hand lane. The wildtype allele (WT) gives a 24kb band, whilst the targeted allele gives a band of approximately 10.4kb. Southern blot of genomic DNA digested with BclI from embryonic stem (ES) cell clones as indicated and wildtype B6 cells, hybridised with P2 are shown, with molecular weight markers in the left hand lane. The wildtype allele (WT) gives a 24kb band, whilst the targeted allele gives a band of approximately 15.3kb. C) PCR genotyping of genomic DNA extracted from mouse pups from a cross between a heterozygous general deleter *βactin-Cre* transgenic mouse and a *Chd8^floX/flox^* mouse. Results from PCR reactions to detect the Cre transgene, distinguish the *Chd8^flox^* and wildtype alleles from each other, and amplify the null allele are shown. Note the loss of the flox allele, with the gain of the null allele in the Cre+ pup (lane 4). D) Quantitative RT-PCR for *Chd8* on mRNA extracted from *Chd8* heterozygous mouse neocortices at E12.5 and P5 and littermate controls using primers spanning the exon3-4 boundary. *Chd8* expression levels in heterozygous mice are significantly reduced to 36% of wildtype controls at E12.5 (**p=0.0059, t=5.346, df=4, student’s t-test, n=3 per genotype) and 48% at P5 (*p=0.0105, t=4.538, df=4, student’s t-test, n=3 per genotype). E) Western blot on lysates from *Chd8* heterozygous E12.5 neocortices and littermate controls. Upper panel: The band for full-length CHD8 (arrow, ∼290kDa) was quantified in F. Note the absence of detectable levels of truncated protein products for CHD8. Lower panel: Western blot for the loading control GAPDH. F) Quantification of CHD8 protein levels normalised to GAPDH as shown in E. CHD8 protein levels in heterozygous mice is 49% compared to wildtype littermates (p=0.156, student’s t-test, n=3 per genotype). G) Quantitative RT-PCR for *Chd8* on mRNA extracted from *Chd8* heterozygous mouse neocortices at E12.5 and littermate controls using primers spanning the exon1-2 boundary (p=0.116, student’s t-test, n=3 per genotype). Samples are the same as in D.

**Supplementary Figure 2:**
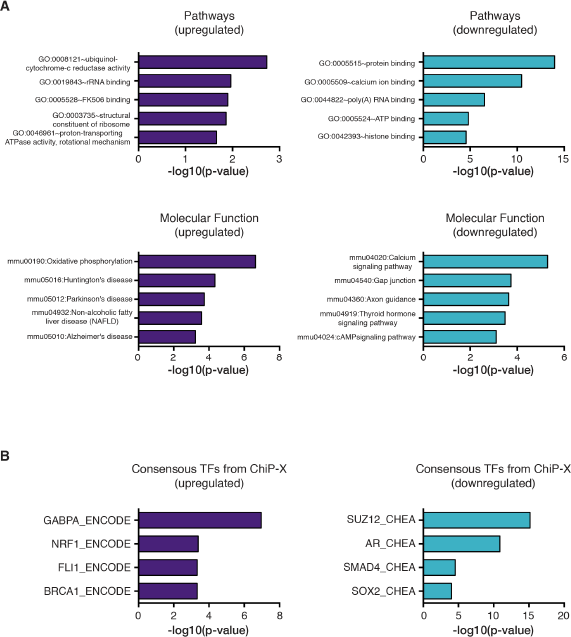
Functional Enrichment Analysis of differentially expressed genes (DEGs) in *Chd8^+/-^* P5 neocortices. A) Results of gene set enrichment analysis using the DAVID knowledgebase on the P5 DEG set (FDR < 0.05). The five most significantly enriched KEGG pathways are shown for upregulated DEGs (top left panel) and downregulated DEGs (top right panel), respectively. The five most significant Gene Ontology terms in the Molecular Function category are shown for up-regulated DEGs (bottom left panel) and downregulated DEGs (bottom right panel). B) DEGs (FDR < 0.05) were subjected to enrichment analysis using the “ENCODE and ChEA Consensus TFs from ChIP-X” database on Enrichr, to identify putative upstream regulatory transcription factors. The four most overrepresented transcription factors are shown for up-regulated (left) and down-regulated (right) DEGs, respectively.

**Supplementary Figure 3:**
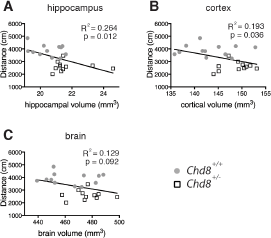
Linear regression analysis of regional brain volumes and activity levels in adult male mice. Activity levels in the open field test were plotted against hippocampal volume (A), cortical volume (B) and brain volume (C) in individual male mice *(Chd8^+/+^,* n=12; *Chd8*^+^", n=l 1). Linear regression analysis: R^2^ =0.264, p=0.012 (hippocampus), R^2^ =0.193, p=0.036 (cortex), R^2^=0.129, p=0.092 (brain).

**Supplementary Figure 4:**
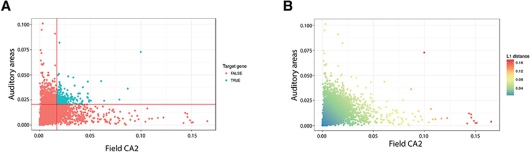
Gene expression enrichment analysis in hippocampus (CA2) and auditory areas. (A) Intersection of genes with high relative expression in the CA2 region and auditory areas. High relative expression was defined as being in the top 20% of genes in a given area. (B) Genes colour coded by their summed relative expression (L1 distance) in CA2 and auditory areas.

**Supplementary Figure 5:**
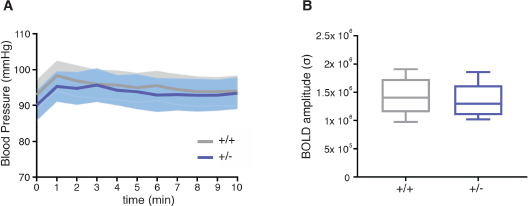
Anaesthesia sensitivity in *Chd8^+/-^* mice and wildtype littermate controls. (A) Continuous monitoring of arterial blood pressure in *Chd8^+/-^* mice and *Chd8^+/+^* littermate controls. Blood pressure does not differ significantly between groups (p=0.79, t=0.274, df=40; student’s t-test). (B) Mean BOLD amplitude in motor cortex of *Chd8^+/-^* mice and *Chd8^+/+^* littermate controls. Mean BOLD amplitude does not differ significantly between groups (p=0.56, t=0.535, df=40; student’s t-test).

## Supplementary Figures

Supplementary Fig. 1: Construction and validation of the *Chd8* conditional and null alleles.

Supplementary Fig. 2: Functional Enrichment Analysis of differentially expressed genes (DEGs) in *Chd8^+/-^* P5 neocortices.

Supplementary Fig. 3: Linear regression analysis of regional brain volumes and activity levels in adult male mice.

Supplementary Fig. 4: Gene expression enrichment analysis in hippocampus (CA2) and auditory areas.

Supplementary Fig. 5: Anaesthesia sensitivity in *Chd8^+/-^* mice and wildtype littermate controls.

## Supplementary Tables

Supplementary Table 1: Absolute and relative volumetric differences in specific brain regions between *Chd8^+/-^* and *Chd8^+/+^* mice as determined by MRI.

Supplementary Table 2: Differentially expressed genes in E12.5 *Chd8^+/-^* neocortices compared to wildtype controls.

Supplementary Table 3: Differentially expressed genes in P5 *Chd8^+/-^* neocortices compared to wildtype controls.

Supplementary Table 4: SFARI ASD genes overlapping with P5 differentially expressed genes.

Supplementary Table 5: Up-regulated Gene Ontology: Biological Processes

Supplementary Table 6: Down-regulated Gene Ontology: Biological Processes

Supplementary Table 7: Up-regulated Gene Ontology: Molecular Function

Supplementary Table 8: Down-regulated Gene Ontology: Molecular Function

Supplementary Table 9 Up-regulated Gene Ontology: Pathways

Supplementary Table 10: Down-regulated Gene Ontology: Pathways

Supplementary Table 11: Gene expression enrichment analysis and Gene Ontology analysis in hippocampal CA2 and auditory areas.

